# Dissecting the midlife crisis: Disentangling social, personality and demographic determinants in social brain anatomy

**DOI:** 10.1101/2020.12.20.423702

**Authors:** Hannah Kiesow, Lucina Q. Uddin, Boris C. Bernhardt, Joseph Kable, Danilo Bzdok

## Abstract

In any stage of life, humans crave social connection with other people. In midlife, transitions in social networks can be related to new leadership roles at work or becoming a caregiver for aging parents. Previous neuroimaging studies have reported that during midlife, especially the medial prefrontal cortex (mPFC) undergoes structural remodeling changes. Social behavior, personality predisposition, and demographic profile all bear intimate relation with the mPFC according to separate literature streams. To integrate these three areas commonly studied in isolation, we explicitly modeled their unique links with brain structure using a fully probabilistic framework. We weighed against each other a rich collection of 40 traits with their interindividual variation in social brain morphology in ~10,000 middle-aged UK Biobank participants (40-69 years at recruitment). Across conducted analyses, household size and daily routine schedules showed several of the largest effects in explaining variation in social brain regions. We revealed male-biased effects in the dorsal mPFC and amygdala for job income, and a female-biased effect in the ventral mPFC for health satisfaction. Our population investigation offers a more complete perspective into how adults at the midlife milestone may navigate life depending on their identity and status.

## Introduction

Humans are inherently social organisms. Already in the first days of life, infants show signs of distress in the absence of social stimulation (Nagy, 2008). As humans grow older, a thirst for social embeddedness with others persists, and may even intensify (Snowden, 2001; Mellor, Stokes et al., 2008). At the midlife point, adults have accumulated decades of social experience that shape how they navigate their daily social encounters. For example, the ability to construct accurate social judgments about other individuals was shown to continuously refine throughout life (Hess and Auman, 2001). Thus, the continuous adaptation of social skills well into adulthood may, in turn, influence other traits that impact how we navigate our social environments, such as personality disposition and demographic standing.

Midlife is a special life period when many milestones have often been reached, such as creating a family and establishing oneself in an occupation (Jaques, 1965). At this point, one’s overall social network size tends to shrink (Charles and Carstensen, 2010; English and Carstensen, 2014). When people have to choose who to spend time with, middle-aged participants were found to prefer familiar over new social interaction partners, compared to younger people (Fredrickson and Carstensen, 1990). Similarly, a behavioral study detailed how often and with whom millions of people communicate on a daily basis (Bhattacharya, Ghosh et al., 2016). The authors confirmed that middle-aged men and women preferentially interact with close family members (Bhattacharya, Ghosh et al., 2016). The consolidation of social networks towards a select core was found to instill a feeling of social embeddedness and increase well-being (Lang, Staudinger et al. 1998; English and Carstensen 2014). Overall, there is an age-related tendency to relatively disengage from social ties at the network periphery in favor of investing in emotionally close others (English and Carstensen, 2014). Hence, midlife can be viewed as a pivotal period in the lifespan when adults transition their focus from exploring new social connections to fostering existing social relationships.

The changes in social network configuration during midlife are probably accompanied by changes in brain architecture. Starting to accelerate in our forties, our brains are known to undergo major structural alterations (Giorgio et al., 2010). A previous voxel-based morphometry study investigated patterns of change in the brain in n=66 participants aged 23 to 81 years (Giorgio et al., 2010). The authors detected increasing reductions in grey matter volume beginning from age 41. The decrease in grey matter structure was especially pronounced in the frontal cortex (Giorgio et al., 2010). This brain region is widely acknowledged for its role in a wide variety of social cognitive processes (Mitchell, 2009). Importantly, during healthy aging, the loss in brain volume is not uniformly distributed across all brain regions (Tisserand, Pruessner et al. 2002; Raz, Lindenberger et al., 2005). Instead, the prefrontal cortex has been emphasized to show the most dramatic and accelerated change in grey matter structure during aging, compared with the rest of the brain (Giorgio et al., 2010; Grieve et al., 2005; Raz et al., 1997; Jernigan et al., 2001; Salat et al., 2004; Good et al., 2001; Tisserand, Pruessner et al., 2002).

Previous brain-imaging research suggests the medial prefrontal cortex (mPFC) serves as a common computational resource for separate domains of social cognition including intrinsic processes related to self-concept or thinking about other people’s mental states (Mitchell, 2009). Similar to how brain structure differs as a function of age, these social cognitive processes may also show characteristic changes across the lifespan. For example, a previous fMRI study investigated differences between younger and older adults in understanding the mental state of other individuals (Moran et al., 2012). In a battery of social-cognitive tasks, the authors found older adults to consistently show deficits compared with younger adults in understanding the thoughts and actions of others, accompanied by isolated reductions in mPFC activation. Moran and colleagues suggest specific involvement of the mPFC in mentalizing skills and that aging is associated with impairments in processing the intentions and internal states of other people (Moran et al., 2012).

In addition to capacities of social interaction, neural activity in the mPFC has also been closely linked to other key domains of everyday life, especially personality and demographics. An individual’s personality is an important source of interindividual variability in approaching everyday life. For example, neuroticism was associated with overall lower well-being during midlife (Abbott, Croudace et al., 2008), as well as grey matter loss in the mPFC (Jackson, Balota et al., 2011). A previous structural brain-imaging study found that ranking high on the personality trait conscientiousness was linked to larger mPFC volumes and less grey matter decline in adults beginning at age 44 (Jackson, Balota et al., 2011). Additionally, extraverted individuals in the later parts of life and individuals whose personality is high on openness to new experience had larger social circles than introverted individuals of the same age (Lang, Staudinger et al., 1998). The collective evidence suggests that personality traits may have an enduring influence on the social behavioral patterns of individuals, and their correlates in brain morphology.

At the broader societal level, interindividual differences in social interaction tendencies also depend on the demographic characteristics of one’s place in society, such as economic resources, occupational prestige and education attainment (Farah, 2017; Muscatell, 2018). Indeed, a brain-imaging study assessed the relationship between socioeconomic disadvantage and grey matter volume (Gianaros et al., 2017). The authors reported smaller grey matter volume in the mPFC for middle-aged adults who experience financial hardship compared with financially stable adults. In addition, financial hardship was also linked to smaller volumes in the hippocampus and amygdala, two limbic regions with direct axonal connections to the mPFC (Butterworth, Cherbuin et al., 2012). Moreover, middle-aged adults with higher education attainment displayed increased amounts of cortical grey matter in the frontal lobe compared with adults with lower levels of education attainment (Kim, Seo et al. 2015). Notably, the protective advantages of high education on cortical thickness were most prominent towards the later midlife years (Kim, Seo et al. 2015). These structural brain-imaging studies exploring different aspects of demographics thus suggest that differences in broader sociocultural experiences may have characteristic imprints in mPFC structure at midlife.

In these many ways, earlier neuroimaging findings highlight the medial prefrontal cortex as a hub that bridges key facets at the individual, interpersonal and societal level. These disparate domains, usually studied in isolation, may reflect common or distinct manifestations in the human social brain. Additionally, many such neuroscience studies have been based on small participant samples as well as samples of student-age (Falk et al., 2013). Therefore, we tailored a probabilistic generative modelling approach to model the extent of similarity and divergence in a rich set of measures from lifestyle domains. Our fully probabilistic modeling framework directly tested against each other 40 traits that track everyday experiences in 10,000 UK Biobank participants in midlife. This modelling tactic allowed us to directly identify the unique contribution of each examined trait to regional volume expressions in the social brain.

## Material and methods

### Population data resource

The UK Biobank initiative (UKBB) is a prospective epidemiology resource that contains a vast portfolio of behavioral and demographic assessments, medical and cognitive measures, and biological samples from a cohort of ~500,000 participants recruited across the United Kingdom (Sudlow, Gallacher et al., 2015). This openly accessible population dataset aims to provide multimodal brain-imaging for ~100,000 individuals to be completed in 2022 (Miller et al., 2016). The present study focused on T1-weighted structural brain magnetic resonance imaging (MRI) from the data release that contained 10,129 individuals, comprising 47.6% males and 52.4% females. The middle-aged cohort was aged between 40 to 69 years at the time of recruitment (mean = 55 years, SD = 7.5 years). All participants were uniformly assessed and brain-scanned at the same scanning facility (i.e., Cheadle). To ensure comparability and reproducibility with other and future UKBB population studies, we used uniform data preprocessing pipelines (Alfaro-Almagro, Jenkinson et al., 2018). The present analyses were conducted under UKBB application number 23827. All participants provided informed consent to participate (http://biobank.ctsu.ox.ac.uk/crystal/field.cgi?id=200).

The present population neuroscience study co-analyzed a total of 40 behavioral indicators provided by the UKBB. The 40 summary measures were divided into three different domains: i) personality (15 items), ii) social (12 items), and iii) demographic (13 items) (cf. Supplementary Table 1). All UKBB participants were administered questions for the particular trait measures (see here for further details: https://www.ukbiobank.ac.uk/). For example, to obtain a measure of the risk-taking trait, participants were asked “Would you describe yourself as someone who takes risks?”. According to the responses given by the UKBB participants for each interview question, participants were evenly split into two groups that reflect the presence or absence of the particular trait: (a) “not a risk-taker” and (b) “risk-taker”. In order to achieve direct comparability of the equal variable encoding in the rich collection of lifestyle indices, all items were represented with binary encoding (Kiesow, Dunbar et al. 2020).

### Brain-imaging preprocessing procedures

The brain-imaging measurements were acquired with a MRI scanner (3T Siemens Skyra) at the same dedicated site (i.e., Cheadle), with the same acquisition protocols and standard Siemens 32-channel radiofrequency receiver head coils. To protect the anonymity of the study participants, brain-imaging data were defaced and any sensitive information from the header was removed. Automated processing and quality control pipelines were deployed (Alfaro-Almagro, Jenkinson et al., 2017). To improve homogeneity of the imaging data, noise was removed by means of 190 sensitivity features. This approach allowed the reliable identification and exclusion of problematic brain scans, such as scans with excessive head motion.

The structural MRI data were acquired as high-resolution T1-weighted images of brain anatomy using a 3D MPRAGE sequence at 1 mm isotropic resolution. Preprocessing included gradient distortion correction, field of view reduction using the Brain Extraction Tool (Smith, 2002) and FLIRT (Jenkinson and Smith 2001; Jenkinson, Bannister et al. 2002), as well as non-linear registration to MNI152 standard space at 1 mm resolution using FNIRT (Andersson, 2007). To avoid unnecessary interpolation, all image transformations were estimated, combined and applied by a single interpolation step. Tissue-type segmentation into cerebrospinal fluid, grey matter and white matter was applied using FAST (FMRIB’s Automated Segmentation Tool, Zhang, Brady et al., 2001) to generate full bias-field-corrected images. In turn, SIENAX (Smith, Zhang et al., 2002) was used to derive volumetric measures normalized for head sizes. The ensuing adjusted volume measurements represented the amount of grey matter corrected for individual brain sizes.

### Social brain atlas definition

Our study benefited from a recently available atlas of the social brain (Alcala-Lopez, Smallwood et al., 2018), which provides a current best estimate of social brain topography in humans. This atlas resulted from quantitatively synthesizing ~4,000 experimental functional MRI studies, involving over ~20,000 participants (Alcala-Lopez, Smallwood et al., 2018). Thirty-six locations of interest were derived in a data-led fashion that were consistently involved in a wide assortment of social and affective tasks (See Supplementary Table 2 for stereotaxic MNI coordinates for each of the social brain atlas regions).

The 36 data-derived social brain regions are connectionally and functionally segregated into four brain networks (cf. Supplementary Table 2): i) a visual-sensory network (fusiform gyrus, posterior superior temporal sulcus, MT/V5), ii) a limbic network (amygdala, ventromedial prefrontal cortex, rostral anterior cingulate cortex, hippocampus, nucleus accumbens), iii) an intermediate network (inferior frontal gyrus, anterior insula, anterior mid-cingulate cortex, cerebellum, supplementary motor area, supramarginal gyrus), and iv) a higher-associative network (dorsomedial prefrontal cortex, frontal pole, posterior mid-cingulate cortex, posterior cingulate cortex, precuneus, temporo-parietal junction, middle-temporal gyrus, temporal pole).

The topographical specificity of the present targeted analyses was thus enhanced by guiding brain volume extraction of the 36 known regions of interest. Neurobiologically interpretable measures of grey matter volume were thus extracted in the ~10,000 participants by summarizing whole brain anatomical maps guided by the topographical compartments of the social brain. In particular, we applied a smoothing filter of 5mm FWHM to the participants’ structural brain maps to homogenize local neuroanatomical differences (Taebi et al., 2020; Kiesow et al., 2020). Next, grey matter volume was extracted in spheres of 5mm diameter around the consensus location from the atlas, averaging the MRI signal across the voxels belonging to a given target region. We would like to note that using a smaller sphere diameter of 2.5mm or a bigger one of 7.5mm yielded virtually identical results. This way of engineering morphological brain features yielded 36 volume brain variables per participant, that is, as many as the total number of social brain regions. Each of the 36 brain volume variables were subsequently z-scored across participants by centering to zero mean and unit-variance scaled to one. These commonly employed estimates of population brain volume variability (Kernbach et al., 2018; Miller et al., 2016) in social brain anatomy served as the basis for all subsequent analysis steps.

All of the regions of interest used in this study are available online for transparency and reuse at the data-sharing platform NeuroVault (http://neurovault.org/collections/2462/).

### Probabilistic multiple regression of social trait variation

In order to explicitly model the unique contribution of the 40 considered lifestyle traits to explaining variation in a given social brain region, we implemented a generative probabilistic multiple regression approach. With this modelling framework, we were able to “learn from data” with traits that dominate in explanatory contribution from three different domains (cf. above; social behavior, personality and demographics). In this way, we were able to directly interrogate the Bayesian posterior uncertainty intervals of trait effects rather than restricting attention to strict categorical differences. Before implementation of the single region probabilistic analyses, we performed a de-confounding procedure on all 36 target region volumes to remove variation due to head size and body mass index. This data cleaning step was performed using nilearn (http://nilearn.github.io/, version 0.6.2) in Python. The probability model with parameters that varied by biological sex were as follows:

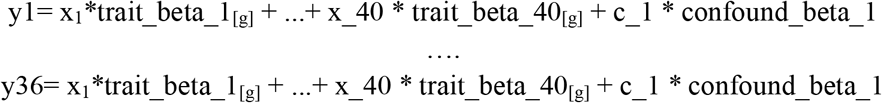

where *x_i_* denotes the behavioral indicator for all 40 considered social lifestyle traits (cf. above), *y_i_* denotes one particular region of the employed social brain atlas (z-scored across participants), and participant group *g* stands for biological sex. Variance that could be explained by the variable *c* of age at recruitment was accounted for as a variable of no interest. In this way, we were able to extract information out of our rich sample as much as possible by borrowing statistical strength between clusters of individuals in our population, to the extent supported by data, through interlocking of their model coefficients.

Approximate posterior inference was achieved by Markov Chain Monte Carlo (MCMC) using PyMC3 in Python (https://github.com/pymc-devs/pymc3, version 3.7), which sampled in a random walk towards the target posterior distribution. In 5,000 draws, the approximate parameter distributions were improved across MCMC steps in the sense of converging to the target distribution. For the partial correlation analysis (cf. below), we carried out the random walk using 10,000 draws to ensure a more stable search of the parameter space. At each step of the MCMC chain, the entire set of parameter values were estimated to be jointly credible given the population data. A range of possible explanations or parameter configurations for the relation between the social lifestyle traits and social brain volume were browsed through by obtaining multiple plausible settings of model parameters that could have led to the observed data. We searched through possible configurations of parameters as an efficient way of exploring the important part of the parameter posterior space. In particular, we dropped the first 4,000 samples from the chain because 1) the chain had probably not yet fully reached stationarity and 2) this step reduced dependence on the starting parameter values. For the partial correlation analysis, we dropped the first 9,000 samples from the chain. Proper convergence was assessed by ensuring the R-hat metrics stayed below 1.02 (Gelman, Carlin et al., 2014).

### Partial correlation analysis

In a subanalysis of our study, we performed a preceding partial correlation step on the 36 region volumes of the social brain atlas using a linear de-correlation procedure. For one specific region among the 36 regions, this complementary analysis partialed out any volume covariation with any of the other respective 35 social brain regions, before carrying out the actual probabilistic regression of interest (cf. above). In this way, we were able to provide a different perspective on how the 40 candidate traits relate to social brain volume in a population cohort. In this variant of our analysis, we interrogated social brain-trait associations after disregarding volume variation shared between any pair of social brain regions.

### Replication analysis

To determine if our probabilistic population results generalize to independent data, we implemented the same data analysis pipelines (cf. above) in new, independent participant samples. First, we randomly split our original n=9,939 brain-imaging UKBB participant dataset into two datasets. The discovery set contained n=4,969 participants, and the replication set had n=4,970 participants. The main region-by-region probabilistic regression analysis was implemented on the discovery set. For the replication analysis, the same probabilistic modelling approach was implemented on the replication set. The replication results yielded fairly good replication of the probabilistic results from the discovery set (Supplementary Table 3). Pearson correlations between the probabilistic model posterior values of the discovery and replication sets display the robustness of our probabilistic results. The Pearson correlations between the discovery and replication sets were calculated for the mean of the posterior population trait distributions as well as the 2.5 and 97.5 highest posterior density interval values across all traits for each social brain region. As a next step, we carried out the same partial correlation analysis pipeline (cf. above) that was implemented on the discovery set onto the replication set. Our partial correlation analysis results from the replication set also revealed good replication of the discovery set results (Supplementary Table 4). Pearson correlations between the model posterior values obtained from the discovery set and replication set show moderately good correlations. Thus, the replication analyses in new independent data samples showcase the robustness of our probabilistic modelling approach.

## Results

Our population neuroscience study sought to juxtapose a rich set of 40 lifestyle factors with their manifestations in brain structure in middle age (40 – 69 years at enrollment). The variety of lifestyle indicators offered by the UK Biobank initiative can be placed into three main categories: i) social exchange, ii) personality profile and iii) demographic status. By integrating all of these indicators into the same analysis, we were able to disentangle which specific traits, relative to the other considered candidate traits, contributed most to explaining social brain volume. As follows, we created 36 different probabilistic models, one for each region in the social brain atlas. Henceforth, the term “trait effect” refers to the marginal posterior parameter distributions of the probabilistic models. It reflects the magnitude, directionality and model uncertainty in the brain association of the analyzed traits (cf. Materials and Methods).

### Social brain midline: Overview of key findings

During midlife, regions of the prefrontal cortex were previously shown to undergo structural changes (Giorgio et al., 2010; Grieve et al., 2005; Raz et al., 1997; Jernigan et al., 2001; Salat et al., 2004; Good et al., 2001; Tisserand, Pruessner et al., 2002). We paid special attention to these regions in the prefrontal cortex because they are also known to be intimately involved in social cognition findings (ex. Mitchell, 2009), including broader social aspects that may bear relation to personality and demographics. In addition, previous research has shown that the mPFC has direct connectivity inputs to several limbic brain regions, such as the amygdala and hippocampus (Bzdok et al., 2012; Aggleton et al., 2015) from the medial-temporal limbic system (Mather, 2016).

In particular, our study aimed to disentangle indicators of social dynamics from trait effects linked to personality features and demographic status within the mPFC and its limbic inputs. Our results revealed that several traits related to the richness of one’s regular social interactions contributed to explaining social brain volume in the mPFC and its limbic connections (Fig. 1).

**Fig. 1.**
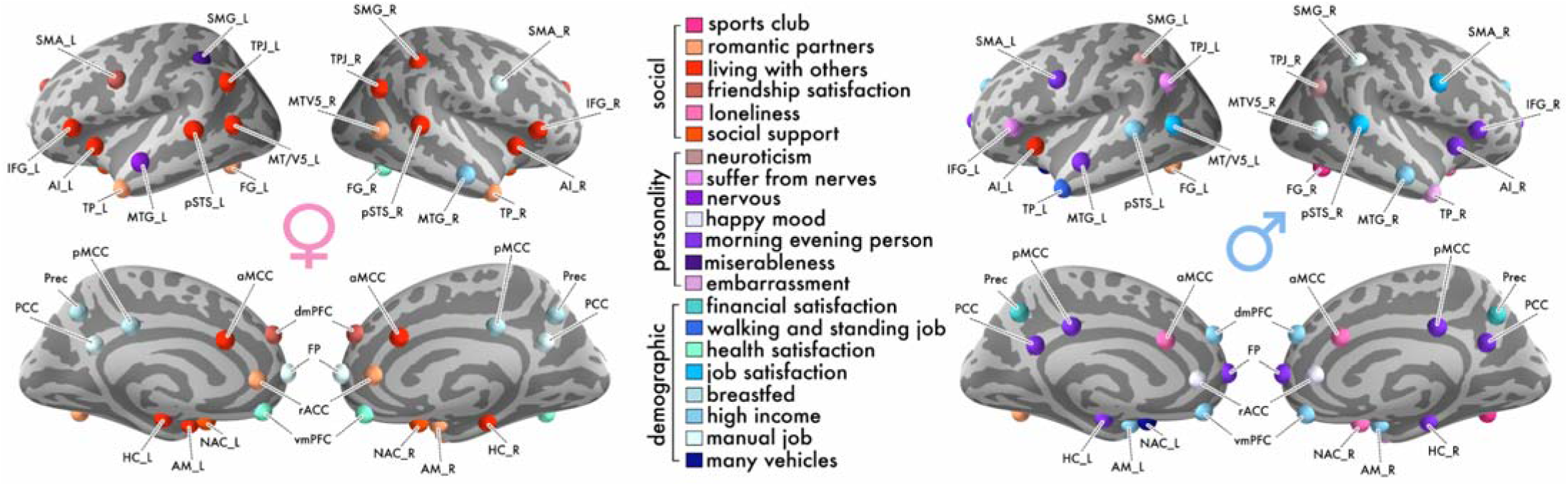
Social brain anatomy is preferentially related to social traits in women, and to personality and demographic traits in men. Extending previous social neuroscience studies, the richness of the UK Biobank resource allowed uniquely isolating partial correlations, which prevail over a wide variety of competing explanatory factors. A generative probabilistic model was applied to each of the 36 social brain regions in our population sample of middle-aged adults. These region-by-region analyses revealed numerous dominant specific (i.e., partial) trait associations in social brain structure. Region colors indicate which individual traits have the largest magnitude (i.e., strongest positive or negative association with region volume) in explaining its regional grey matter volume, relative to the other 39 out of 40 total examined traits. Red colors indicate markers in the social trait category, purple colors of the personality trait category and blue colors of the demographic trait category. Sharing one’s home with other individuals was the single most frequent trait association to show the largest magnitude in explaining social brain volume for women. The personality trait of being a morning person was the most common trait to show the strongest trait association in social brain structure for men. Dominant trait associations from the partial correlation analysis are shown in Supplementary Fig. 1.

For example, sharing the home environment with other individuals was a top contributor to explaining grey matter volume in the limbic AM_L and bilateral HC (see Supplementary Table 2 for abbreviation list; women: AM_L: mean of the population trait posterior distribution = 0.035, highest density interval of the population trait posterior distribution covering 95% model uncertainty (HPDI) = 0.005 – 0.069; HC_L: posterior mean = 0.040, HPDI = 0.012 – 0.072; HC_R: posterior mean = 0.056, HPDI = 0.019 – 0.093). Parallelling these dominant trait findings on household size, markers related to interaction with close social partners, such as friends, showed unique trait associations in our middle-aged population cohort, relative to the other 39 competing traits. For example, feeling satisfied with one’s friendships was a top trait association in the higher associative dmPFC region (women: posterior mean = 0.039, HPDI = 0.000 – 0.078). Moreover, we also observed that the lifetime number of romantic partners showed a dominant trait association in the AM_R (women: posterior mean = 0.031, HPDI = 0.009 – 0.056).

In addition to traits characterizing aspects of one’s social lifestyle, we also found personality trait effects in regions of our social brain atlas (Fig. 1). Specifically, being a morning versus evening person showed the largest magnitude in explaining grey matter volume in the FP and bilateral HC, compared to the other candidate traits (men: FP: posterior mean = 0.052, 95% HPDI = 0.002 – 0.104; HC_L: posterior mean = 0.051, HPDI = 0.020 – 0.084; HC_R: posterior mean = 0.054, HPDI = 0.022 – 0.087).

Regarding traits capturing the societal level, we observed several demographic traits to show unique trait effects in midline social brain regions and limbic inputs (Figs. 1 and 2). Our Bayesian posterior distributions showed that earning a high yearly income explained the biggest fraction of region volume in the vmPFC and bilateral AM, compared to the other considered traits (men: vmPFC: posterior mean = 0.053, 95% HPDI = 0.011 – 0.100; AM_L: posterior mean = 0.060, HPDI = 0.022 – 0.104; AM_R: posterior mean = 0.048, HPDI = 0.017 – 0.084). In the vmPFC, a similarly strong trait effect was observed (Fig. 2). Health satisfaction, a trait linked to SES, was the top contributor to vmPFC grey matter volume compared to the other examined traits (women: posterior mean = 0.067, HPDI = 0.030 – 0.105). In addition, working a manual, as opposed to a knowledge-based job, showed a dominant trait effect in the FP (women: posterior mean = 0.059, HPDI = 0.007 – 0.115). Taken together, our probabilistic evidence revealed interindividual variability in trait variation that is mostly linked to social experience and demographics in social brain midline and limbic regions.

**Fig. 2.**
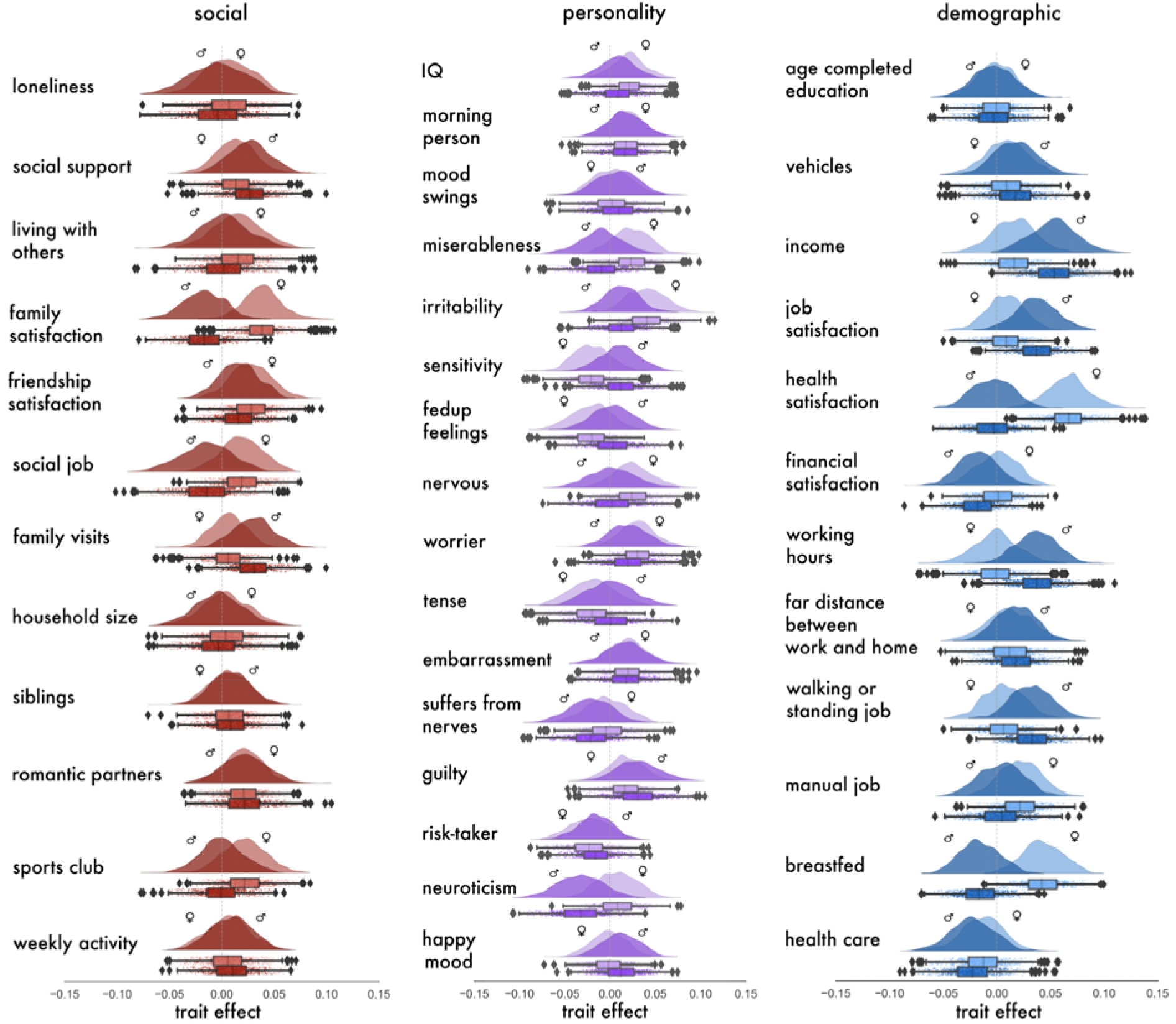
vmPFC anatomy reveals social lifestyle markers of social, personality, and demographic traits to show sex-differentiated population effects. During midlife, regions of the mPFC have been found to show an accelerated decline in grey matter structure (cf. introduction). For the sake of illustration, the marginal posterior population distributions from our vmPFC analysis are depicted using raincloud plots (1 of the 36 region-by-region analyses that we have conducted on the UK Biobank). We reveal to what extent vmPFC region volume is specifically explained by the 40 examined lifestyle traits at the population level. The half violin plots show the posterior parameter distributions of the specific contributions of each single trait to vmPFC volume, not explained by the other traits, in middle-aged men and women. The boxplots and scatterplots underneath depict the probabilistic parameter guesses that together form the marginal posterior distributions. For middle-aged women, satisfaction with health contributed most to explaining vmPFC region volume. In contrast, for middle-aged men, earning a higher job income explained most vmPFC grey matter volume.

Moreover, the reiteration of our analyses based on partial region volumes of the social brain (cf. Materials and Methods) shed a different light on the dominant trait associations in almost all of the social brain midline and limbic regions. The partial correlation analysis also revealed a richer variety of dominant trait effects across atlas regions, mostly in the category of demographic traits (Supplementary Fig. 1; cf. Supplementary information for a full description of the partial correlation analysis results). For example, having an occupation that requires working more than 40 hours a week showed a dominant trait association in the vmPFC (men: posterior mean = 0.005, 95% HPDI = −0.006 – 0.016). However, completing full-time education, another trait indexing SES, showed dominant trait associations in the FP and limbic AM (men: AM_R: posterior mean = 0.009, HPDI = −0.002 – 0.020; women: FP: posterior mean = 0.039, HPDI = 0.008 – 0.075). Taken together, the partial correlation analysis findings for the midline social brain regions revealed a wider assortment of dominant trait associations mostly linked to demographic profile.

### Social markers: Household size is consistently linked to social brain structure during midlife

Our fully probabilistic region-by-region analyses revealed at the interpersonal level that social markers associated with social network composition and quality of interpersonal exchange showed consistently dominant trait effects across social brain regions (Fig. 1). In our middle-aged population cohort, sharing the home environment with other individuals was the most common trait to explain the most region volume in about half of our 36 region analyses.

Living with others, as opposed to living alone, showed dominant trait effects in regions of the higher associative network of our social brain atlas, such as the bilateral TPJ (women: TPJ_L: posterior mean = 0.070, 95% HPDI = 0.015 – 0.127; TPJ_R: posterior mean = 0.059, HPDI = 0.010 – 0.109). Compared to the other candidate traits, sharing the home environment with other individuals also explained the largest fraction of volume variation in regions of the intermediate network, including the bilateral AI and bilateral IFG (men: AI_L: posterior mean = −0.045, HPDI = −0.107 – 0.011; women: AI_L: posterior mean = 0.056, HPDI = 0.012 – 0.101; AI_R: posterior mean = 0.058, HPDI = 0.024 – 0.100; aMCC: posterior mean = 0.070, HPDI = 0.021 – 0.120; IFG_L: posterior mean = 0.039, HPDI = − 0.005 – 0.083; IFG_R: posterior mean = 0.065, HPDI = 0.014 – 0.118; SMG_R: posterior mean = 0.040, HPDI = −0.005 – 0.087). Furthermore, this social trait showed dominant population trait effects in regions of the visual sensory network of the social brain, including the bilateral pSTS and MT/V5_L (women: MT/V5_L: posterior mean = 0.050, HPDI = 0.004 – 0.097; pSTS_L: posterior mean = 0.058, HPDI = 0.010 – 0.110; pSTS_R: posterior mean = 0.049, HPDI = 0.000 – 0.096).

Additionally, social markers related to close interpersonal relationships also revealed large magnitudes in explaining grey matter volume in regions of the visual sensory, limbic, and higher associative networks. In particular, the lifetime number of romantic partners showed dominant trait effects in several visual sensory, limbic and higher associative social brain regions including the FG, rACC and TP (women: FG_L: posterior mean = 0.032, 95% HPDI = 0.005 – 0.060; MTV5_R: posterior mean = 0.054, HPDI = 0.012 – 0.094; TP_L: posterior mean = 0.037, HPDI = 0.010 – 0.067; TP_R: posterior mean = 0.033, HPDI = 0.000 – 0.065; rACC posterior mean = 0.057, HPDI = 0.018 – 0.099; men: FG_L: posterior mean = 0.048, HPDI = 0.009 – 0.086). This pattern of trait effects linked to social interaction with close family and friends was also observed in a region of the visual sensory network. Specifically, being a member of a sports club, a social trait related to social group involvement, showed the largest trait effect in the FG_R (men: posterior mean = 0.045, HPDI = 0.002 – 0.086).

In interindividual differences of social support, regular exchange with emotionally close others explained the most region volume in the bilateral NAC, compared to the other analyzed traits (women: NAC_L: posterior mean = 0.076, 95% HPDI = 0.018 – 0.133; NAC_R: posterior mean = 0.040, HPDI = −0.010 – 0.091). In a related social marker indexing the quality of close relationships, feelings of loneliness showed a unique trait association in several limbic and intermediate network brain regions, including the reward-related NAC (men: NAC_R: posterior mean = −0.051, HPDI = −0.128 – 0.020; aMCC: posterior mean = − 0.051, HPDI = −0.114 – 0.011). Collectively, dimensions on the strength of social closeness to close family and friends were associated most with grey matter volume in a majority of social brain regions. Notably, sharing a home with other individuals was the most frequently observed dominant trait association in our midlife population.

Our analysis based on partial volume correlations revealed dominant trait associations similar to that of the main analysis (cf. Supplementary Fig. 1; cf. Supplementary information for a full description of the partial correlation analysis results). For example, having a job that requires much social interaction was a top frequent trait to contribute to social brain volume in regions of the visual sensory, intermediate and limbic brain networks (women: FG_R: posterior mean = −0.015, 95% HPDI = −0.036 – 0.004; IFG_L: posterior mean = −0.004, HPDI = −0.020 – 0.008; NAC_R: posterior mean = −0.008, HPDI = −0.025 – 0.007; SMA_L: posterior mean = −0.010, HPDI = −0.031 – 0.009). Together, our social trait findings from the partial correlation analysis identified a similar set of dominant trait associations to that of the main analysis.

### Personality markers: Top brain associations are associated with daily routine and well-being

Focusing on trait effects from the personality category, markers associated with daily routines and psychological well-being showed large magnitudes in explaining grey matter volume (Fig. 1). In general, the personality trait of being a morning versus evening person explained the most region volume in 11 different social brain regions, compared to the other candidate traits (Fig. 1). This biorhythm trait showed dominant trait effects in regions of our intermediate network of the social brain atlas, including the AI_R (men: posterior mean = 0.050, 95% HPDI = 0.015 – 0.089), bilateral CB (men: CB_L: posterior mean = 0.069, HPDI = 0.016 – 0.123; CB_R: posterior mean = 0.083, HPDI = 0.028 – 0.140), SMA_L (men: posterior mean = 0.056, HPDI = 0.014 – 0.101), and IFG_R (men: posterior mean = 0.058, HPDI = 0.015 – 0.106). Morning versus evening chronotype also contributed to grey matter volume in regions of the higher associative network, such as the MTG_L (men: posterior mean = 0.043, HPDI = 0.006 – 0.081), PCC (men: posterior mean = 0.051, HPDI = 0.013 – 0.088), and pMCC (men: posterior mean = 0.044, HPDI = 0.012 – 0.075).

Additionally, our region-by-region analyses identified several personality traits related to long-term well-being to show dominant trait effects in several social brain regions belonging to the limbic, intermediate and higher associative networks. For example, for our middle-aged participants, having a happy mood showed the largest trait effect in the limbic rACC (men: posterior mean = 0.059, 95% HPDI = 0.017 – 0.100). However, neuroticism, a personality trait, explained the most variation in several intermediate and higher associative network regions, including the TPJ_R, compared to the other analyzed traits (men: TPJ_R: posterior mean = −0.044, HPDI = −0.106 – 0.015; SMG_L: posterior mean = −0.050, HPDI = −0.113 – 0.009). Together, the personality traits associated with daily routine schedules and personal well-being were top traits to explain grey matter volume in a number of social brain regions.

The complementary partial correlation analysis revealed a wider variety of personality traits to explain social brain region volume in our middle-aged population cohort (Supplementary Fig. 1; cf. Supplementary information for a full description of the partial correlation analysis results). For example, the feeling of miserableness revealed dominant trait associations in several intermediate and higher associative social brain regions including the SMG_L (women: posterior mean = 0.014, HPDI = −0.010 – 0.041) and TP_R (men: posterior mean = 0.006, 95% HPDI = −0.007 – 0.020). Similarly, we observed the risk-taking trait to show dominant trait associations in the limbic HC_R (men: posterior mean = −0.011, HPDI = −0.024 – 0.001) and intermediate SMG_R region (women: posterior mean = 0.008, HPDI = −0.011 – 0.029). Taken together, results from the partial correlation analysis revealed a larger range of personality traits to contribute to social brain grey matter volume, compared to the main analysis.

### Demographic markers: Traits related to income and occupation are associated with limbic and higher associative brain regions during midlife

At the broader societal level, we observed a variety of demographic traits related to social status and occupation to explain social brain grey matter volume in our middle-aged population sample (Fig. 1). In particular, earning a high yearly wage showed dominant trait effects in several higher associative and visual sensory regions of our social brain atlas (men: MTG_R: posterior mean = 0.069, 95% HPDI = 0.027 – 0.111; pSTS_L: posterior mean = 0.065, HPDI = 0.016 – 0.113; women: MTG_R: posterior mean = 0.047, HPDI = 0.006 – 0.090).

Our region-by-region probabilistic results further revealed granularity in aspects related to one’s occupational environment. For example, working a manual job explained the most variation in four social brain regions of the visual sensory, intermediate and higher associative brain networks (men: MTV5_R: posterior mean = 0.042, 95% HPDI = −0.007 – 0.091; SMG_R: posterior mean = 0.058, HPDI = 0.011 – 0.108; women: PCC: posterior mean = 0.042, HPDI = 0.009 – 0.074; SMA_R: posterior mean = 0.070, HPDI = 0.025 – 0.120). Similarly, having a job that requires walking or standing for most of the workday showed a dominant trait effect in the higher associative TP_L (men: posterior mean = 0.045, HPDI = 0.009 – 0.078). In the context of the work environment, feeling satisfaction with one’s occupation contributed to explaining region volume in regions of the visual sensory and intermediate networks (men: MTV5_L: posterior mean = 0.033, HPDI = −0.003 – 0.075; SMA_R: posterior mean = 0.042, HPDI = −0.002 – 0.086; pSTS_R: posterior mean = 0.059, HPDI = 0.018 – 0.102). Together, our dominant demographic trait findings at the general societal level revealed that during midlife, several aspects of one’s job environment contributed most to explaining gray matter volume variation in a variety of social brain regions.

Results from the partial correlation analysis revealed that demographic traits related to occupation were also the top contributing traits in a number of social brain regions (cf. Supplementary Fig. 1; cf. Supplementary information for a full description of the partial correlation analysis results). For example, feeling satisfied with one’s occupation was the most frequent trait to contribute to social brain volume in several limbic, intermediate and higher associative network brain regions (men: AI_R: posterior mean = −0.011, 95% HPDI = −0.032 – 0.002; IFG_R: posterior mean = 0.010, HPDI = −0.011 – 0.034; NAC_L: posterior mean = −0.025, HPDI = −0.052 – 0.000; NAC_R: posterior mean = 0.007, HPDI = −0.006 – 0.021; Prec: posterior mean = −0.007, HPDI = −0.023 – 0.008; women: pSTS_R: posterior mean = 0.010, HPDI = −0.004 – 0.030). Similarly, the partial correlation analysis showed that working more than 40 hours a week contributed to social brain grey matter volume (men: CB_L: posterior mean = −0.007, HPDI = −0.020 – 0.004; MTG_R: posterior mean = 0.007, HPDI = −0.008 – 0.024; MT/V5_L: posterior mean = −0.010, HPDI = −0.032 – 0.009; vmPFC: posterior mean = 0.005, HPDI = −0.006 – 0.016). Taken together, the demographic trait findings from the partial correlation analysis revealed that the top trait associations most associated with social brain grey matter volume encompassed traits related to occupation.

In summary, our region-by-region analyses on a diverse selection of social, personality and demographic traits together showed manifestations in social brain grey matter structure in a population cohort of adults at the midlife turning point. Compared to the social and demographic domains, traits from the personality category showed less dominant trait associations for our middle-aged participants. Instead, markers indexing social exchange at the interpersonal and broader societal level contributed more to explaining volume variation in social brain regions during midlife.

### Age drives trait associations dominant in mPFC and limbic regions during midlife

Our analyses also revealed considerable contributions of age to jointly explain grey matter variation with other lifestyle traits (Fig. 3; cf. Supplementary Fig. 2 for results from the partial correlation analysis). Furthermore, most of the joint age-trait effects in the medial prefrontal cortex and its interaction partners from the limbic system were manifested differently in our sample of middle-aged men and women.

**Fig. 3.**
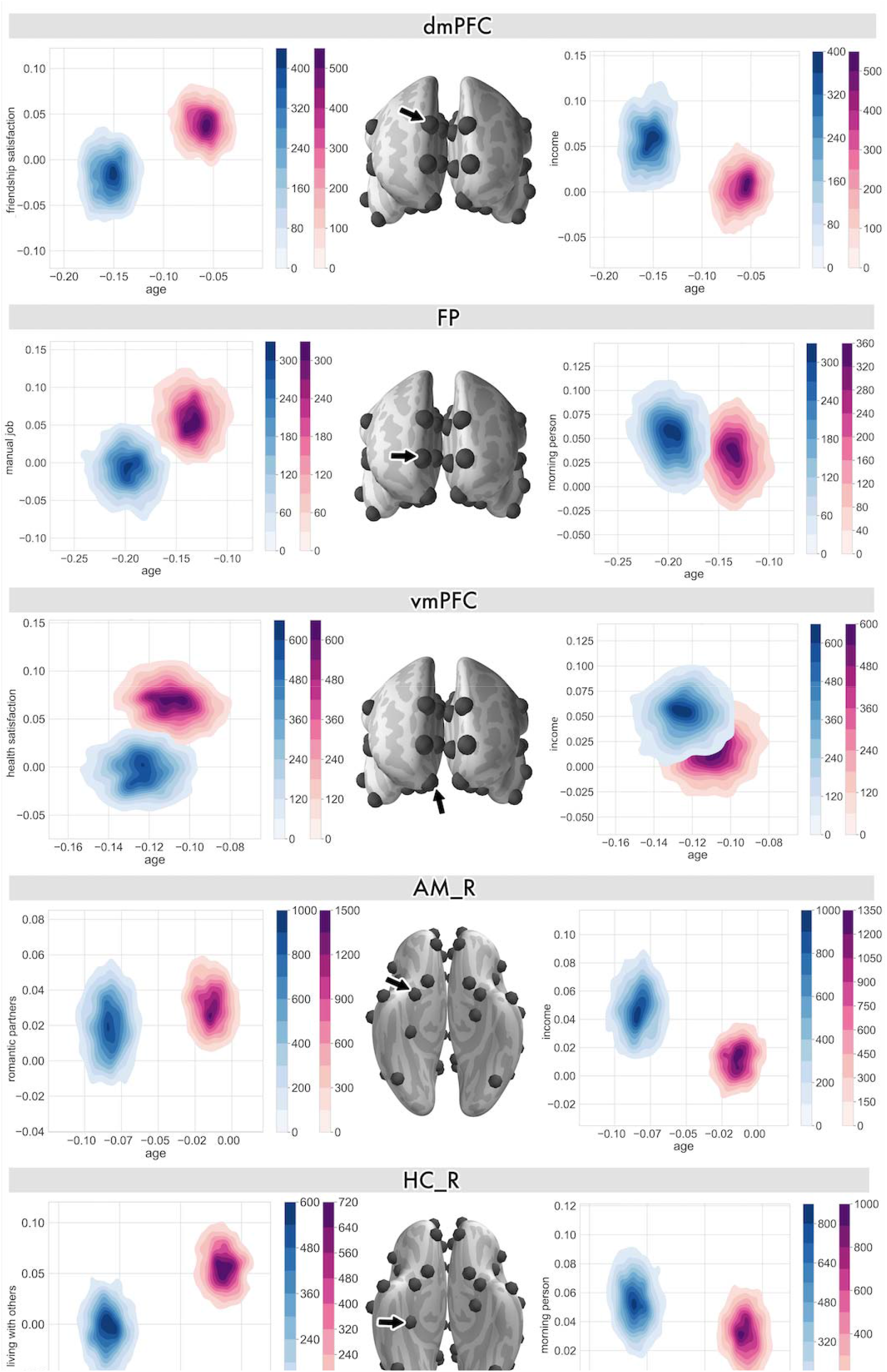
Participant age drives how dominant lifestyle traits are linked to midline social brain regions. The co-relationship between age and a trait association is quantified by the joint posterior parameter distribution for one particular social brain region (black arrow) for men (blue) and women (pink). For this summary visualization, the traits were picked based on the top effects in the full analysis (cf. Fig. 1). The left column shows the top trait associations for women and the right column shows the top traits for men. Middle-aged men and women showed diverging age-trait associations in dmPFC volume in the context of social interaction quality and social status. However, in the context of two SES measures, men and women were more similar in age-trait associations with vmPFC volume. The joint posteriors of trait associations in the FP revealed socioeconomic status as measured by job type and personality to show incongruent population parameter distributions. The limbic AM and HC regions additionally showed non-overlapping posterior distributions between middle-aged individuals for social network size and social lifestyle. Supplementary Fig. 2 showcases the age-trait associations in the midline and limbic regions from the partial correlation analysis.

For example, in the dmPFC, age and high friendship satisfaction (the top trait association in the dmPFC for women) were together associated with largely divergent manifestations of dmPFC region volume for men and women across midlife (Fig. 3). Similarly, age and earning a higher income (the top trait association in the dmPFC for men) was differentially linked to dmPFC region volume for men and women. However, as a function of age, working a manual job was related to FP variation. For women, working a manual job was the top FP trait association (cf. Fig. 1). Similarly, age and the personality disposition of being a morning person showed only slightly overlapping posterior parameter distributions in the FP for men and women (Fig. 3). For the top female trait association in the vmPFC, high health satisfaction and age jointly explained region variation with incongruent posterior distributions for men and women. In conjunction with age, having a high yearly job income (the top male trait association in the vmPFC) showed overlapping model posteriors in explaining vmPFC volume variation for men and women.

Furthermore, we also observed age to jointly drive the dominant trait effects in several limbic regions of the social brain that are known to have connections to the mPFC. For example, age and the number of lifetime romantic partners (the top female trait association in the amygdala) showed largely divergent posterior distributions for men and women. Moreover, high job income (the top male trait association) showed large posterior divergences in explaining grey matter volume in the AM_R for men and women. In the memory-related region of the social brain, age and sharing a home with other individuals influenced hippocampal architecture differently for men and women. In conjunction with age, morning chronotype (the top male trait association in the HC_R) showed an opposite pattern in hippocampal region volume for men and women in our population sample.

Together, the joint age-trait posterior distributions revealed mostly divergent manifestations of grey matter volume in midline and limbic brain regions for men and women. For women, these population volume effects were more apparent in sociodemographic and social indicators, as a function of age. For men, age and sociodemographic and personality indicators were more prominent.

### Network-by-network summary: Specific social brain regions are better explained by the collective traits than others

In each of the four networks of the social brain (cf. Materials and Methods), we calculated posterior predictive checks for each region model. This diagnostic assessment is based on simulation of new replicated data using our previously inferred probabilistic models (Gelman, Carlin et al., 2014). The simulated model outcome can then be compared to the actual observed outcomes to get a sense of our already estimated model parameter distributions. For each of the 36 examined regions from the social brain atlas, we computed the amount of explained variance from the posterior predictive checks (coefficient of determination, R^2^) (Fig. 4). In this way, we interrogated which of the four networks were best explained from the 40 examined lifestyle traits.

**Fig. 4.**
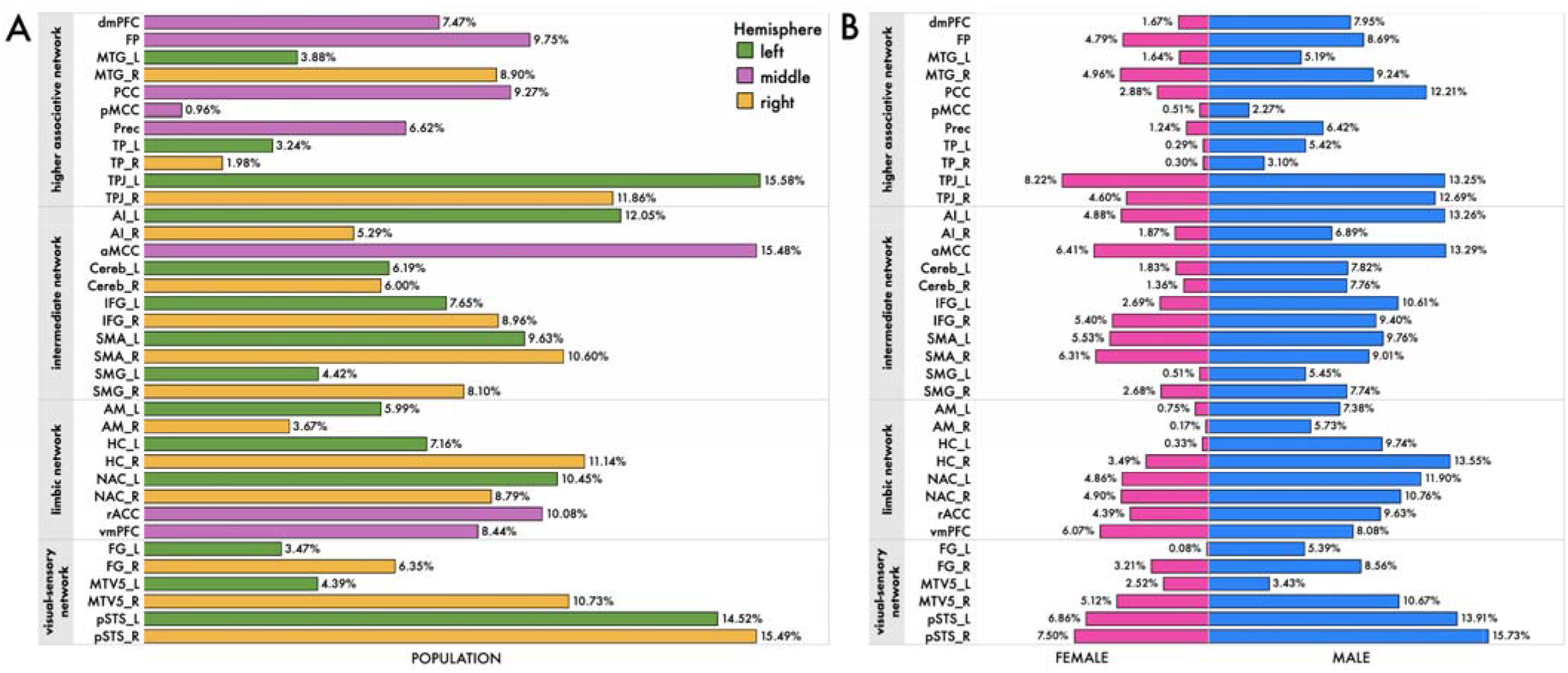
Total amount of explained variance in lifestyle trait associations differs across social brain networks. In each of the four networks of our social brain atlas (cf. Materials and Methods), we computed posterior predictive checks for every region analysis. These model-based simulations of replicated data were then compared to the actually observed data (Gelman, Carlin et al., 2014) to compute the overall explained variance (coefficient of determination, R^2^) for A) the whole population cohort and B) men and women, separately. The collective population-level results suggest that in each of the four subnetworks of the social brain atlas, at least one region successfully explained >10% of region volume in our middle-aged participant cohort.

The total explained variance was highest for the bilateral pSTS (pSTS_R: ~16%; pSTS_L: ~15%), two regions of the visual-sensory network (Fig. 4). In the intermediate network, aMCC and AI_L volumes showed the highest total explained variance (aMCC: ~16%; AI_L: ~12%). Instead, the TPJ_L (~16%) and TPJ_R (~12%) were the most explanatory parts of the higher associative network. In the limbic network, the HC_R (~11%) and NAC_L (~11%) showed the best R^2^ scores. Together, the collective population results suggest that in each of the networks of the social brain atlas, at least one region was able to explain >10% of region volume in our population sample.

### Health satisfaction and income show specific trait effects in midline and limbic regions

Finally, we directly quantified the extent of sex differentiation in our brain-trait associations by calculating the difference beween the marginal posterior distributions. We carried out the subtraction (female – male) of the model posterior parameter distributions for each trait in each analysis of a social brain region. The obtained difference contrasts of the posterior parameter distributions could reveal relatively more male- or more female-driven effects for a specific trait. In the medial prefrontal regions and its limbic input regions, we observed more male-driven population trait effects (Fig. 5; cf. Supplementary Fig. 3 for results from the partial correlation analysis). However, results in the FP showed more female-specific trait effects. Compared to the other midline social brain regions, the single-trait posterior distributions for the FP also showed much more uncertainty in the difference contrasts of the model posteriors.

**Fig. 5.**
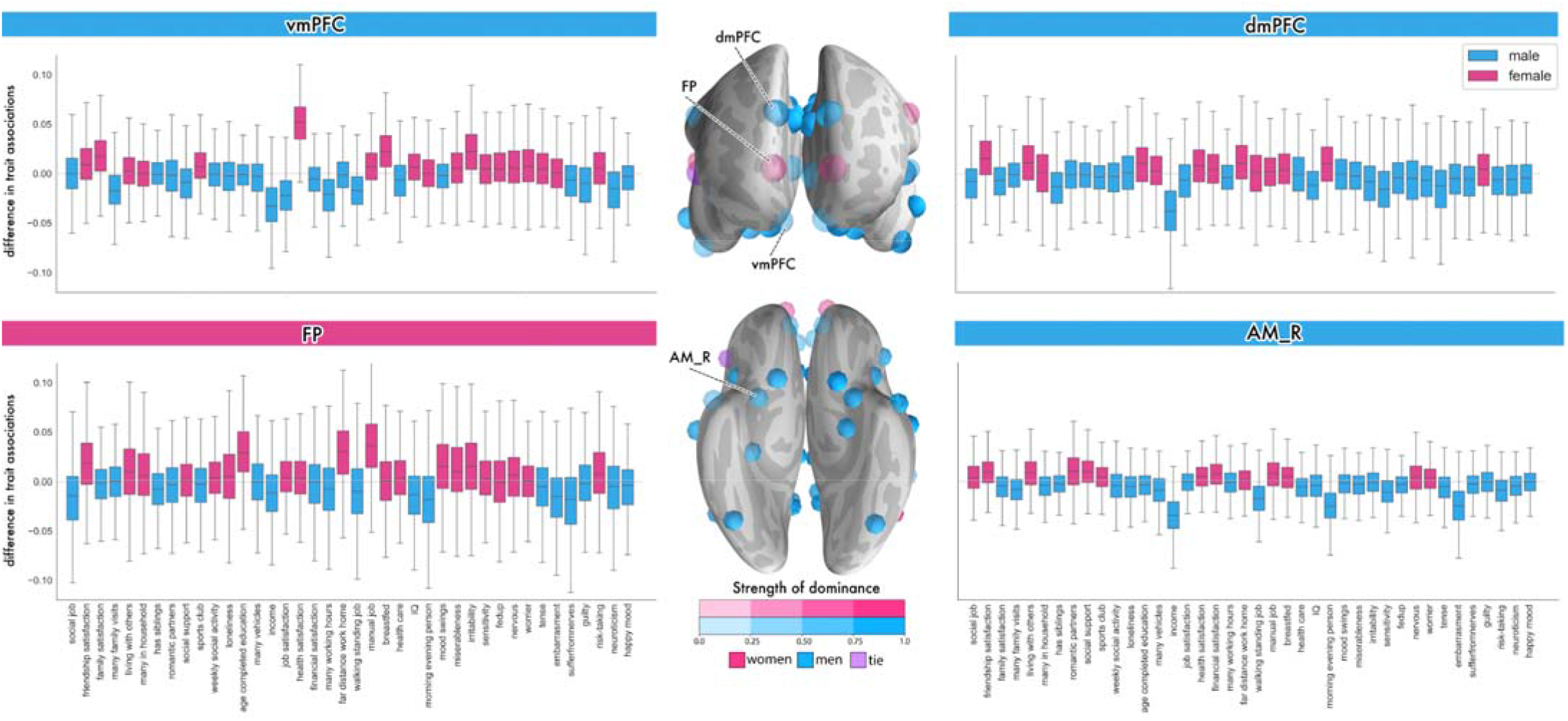
Degree of sex bias in lifestyle trait associations in the social brain midline. Left/Right: Boxplots depict the difference contrasts of the marginal posterior population distributions from each sex (female – male) in each of the 40 examined traits for the social brain midline. As such, posterior distribution means above zero indicate a relatively female-biased effect for that specific trait association (pink). For values below zero, there is a relatively male-biased effect for that specific trait (blue). Middle: As a summary visualization, for each brain region, we counted across the 40 trait associations how many were biased predominantly towards males (blue) or more females (pink). Purple shows an equal number of male- and female-biased trait associations. Transparency indicates the strength of the sex effect. Overall, the population-level findings revealed a male bias in volume effects in almost all examined midline and limbic regions. In the dmPFC and AM_R, yearly job income showed a stronger effect in men compared to women. However, women drive the trait associations in the FP of the higher-associative social brain, especially with regards to several demographic traits, such as the age of full-time education completion and working a manual job (cf. Supplementary Fig. 3 for sex differentiation in lifestyle trait associations from the partial correlation analysis).

In the vmPFC, a large female-specific trait effect became apparent. Health satisfaction contributed more to explaining volume variation in the vmPFC for women than for men. Further, in the dmPFC and AM_R regions, we observed a few specific, strong male-biased population trait effects. Notably, in the dmPFC and AM_R, earning a higher job income showed a much larger effect for men, compared to women. In addition, the feeling of embarrassment and the personality trait of being a morning person showed larger trait effects in the AM_R for men compared with women. Together, the findings from the sex contrast analyses revealed degrees of sex differentiation in population associations in the limbic and higher associative regions across trait domains.

## Discussion

During midlife, many aspects of the social environment and lifestyle are subject to change (Jaques, 1965). Previous neuroimaging research has shown that during this period of life, the brain also undergoes concomitant structural changes, especially in its midline regions (Giorgio et al., 2010). Using fully probabilistic modelling, our study interrogated social brain architecture through the prism of 40 indicators of social, personality, and demographic profile in ~10,000 UKBiobank participants. Many previous social neuroscience studies were limited in juxtaposing a wide breadth of indicators, partly because of the scarcity of datasets that provide deep phenotyping with diverse groups of behavioral assessments. Here, we submitted such a rich collection of lifestyle indicators covering the individual, interpersonal and broader societal level into one coherent analysis framework.

Previous neuroimaging studies have reported that the prefrontal cortex shows an accelerated structural decline with increasing age compared with any other part of the brain (e.g., Giorgio et al., 2010; Grieve et al., 2005; Raz et al., 1997; Jernigan et al., 2001; Salat et al., 2004; Good et al., 2001; Tisserand, Pruessner et al., 2002). The vulnerability for prefrontal atrophy may relate to the notion that regions of the brain that mature later in development may be more susceptible to structural changes (Grieve et al., 2005). For example, a structural MRI study investigated cortical atrophy in young (18-31 years), middle (41-57 years) and old (60-93) age groups (Salat et al., 2004). The authors found global cortical thinning in the brain starting at midlife. This previous study also showed that cortical atrophy is the most pronounced in the prefrontal cortex compared with the rest of the brain. Another structural MRI study reported age-related cortical decline in several regions of the mPFC (Grieve et al., 2005). The authors speculated that loss in grey matter volume may be linked to replacement of volume by cerebral spinal fluid or an increase in white matter.

More generally, the mPFC is increasingly recognized to be a common denominator for disparate fields of neuroscience (Mitchell, 2009), especially social interaction (Moran, Jolly et al., 2012), personality (Jackson, Balota & Head, 2011) and demographics (Muscatell, Morelli et al., 2012). Integrating these separate results point to the mPFC to act as a hub, or a common neurocomputational resource for different processes related to daily experiences and social identity. Our middle-aged population results thus detail and extend previous findings by showcasing in ~10,000 individuals that interindividual variation in grey matter volume in regions of the social brain is linked to key aspects of day-to-day experience and lifestyle determinants. Here, we highlight that at midlife, when the brain and especially its mPFC are known to start to decline, aspects related to social support and social status may contribute most to explaining variation in brain volume.

Elements of daily experiences include interacting with close friends and family members. Previous behavioral studies have highlighted that during midlife, adults selectively interact with emotionally close others (English & Carstensen, 2014). Maintaining close social bonds depends on mentalizing, or understanding the mental states of other individuals (Dunbar, 2018). As part of the higher associative network of the social brain (Alcala-Lopez et al., 2018), the dmPFC is widely acknowledged for its role in such Theory-of-Mind processes (Mitchell, 2009), which involve taking the perspective of other individuals to understand their emotions, beliefs, and motivations (Frith & Frith, 2005). Here, we observed that, comparing 40 traits against each other, friendship satisfaction showed a dominant trait effect in the dmPFC.

A previous behavioral study reported that feeling unsatisfied with personal relationships is associated with the experience of loneliness (Mellor et al., 2008). Loneliness is a subjective perception that can have wide ranging consequences, including decreased mental and psychological well-being (Bzdok & Dunbar, 2020), and even increased mortality (Holt-Lunstad et al., 2015). Indeed, a cohort study on loneliness in ~900 middle-aged participants (Nersesian et al., 2018) found that loneliness was linked to higher levels of stress and systemic inflammation. The authors suggest that these observations are in line with poor health outcomes, with increased risk for morbidity and mortality. This constellation of findings adds to the idea that the amount of regular investments in social networks are closely linked to our dmPFC measurements.

Consistent with the perspective that adults characteristically place their social investments on people already in their social circles during midlife, we found markers of social closeness to also relate to the AM and HC − two limbic brain regions with close anatomical connections to the mPFC, as evidenced in humans and monkeys (Bzdok et al., 2012; Aggleton et al., 2015). Gauging 40 candidate traits, our results singled out the lifetime number of romantic partners as a top trait to explain variation in AM volume. Consistently, previous structural brain-imaging studies have also linked the AM to variation in social group size in several age groups (Bickart, Wright et al., 2011; Kiesow, Dunbar et al., 2020). In particular, a relationship was reported between larger grey matter volume in the AM and increasing size and complexity of social networks (Bickart, Wright et al., 2011). Similarly, social support and household size, two markers intimately linked to social network size and complexity, were found to be associated with AM volume in a previous population neuroscience study (Kiesow, Dunbar et al. 2020).

In a similar vein, our results from a middle-aged cohort revealed that sharing the home environment with others showed a dominant trait effect in the HC, a limbic region with intimate functional and structural ties to the mPFC (Bzdok, Langner et al., 2013; Cohen, 2011). Part of the medial temporal lobe, grey matter hippocampal volume has also been observed to be related to differences in social network size (Kanai, Bahrami et al., 2012). Furthermore, a previous behavioral study found that middle-aged adults living alone less often reported feeling satisfied with their personal relationships, suggesting an unmet need for belongingness (Mellor et al., 2008). Together, these considerations hint at a particularly strong role of the mPFC and limbic regions in investing in close relationships with friends and family.

In addition to understanding the mental states of other individuals, the mPFC has also been found to be implicated in other social cognition processes (Mitchell, 2009). Social status is an abstract construct of individuals’ standing in society compared to others (Festinger, 1954; Zink et al., 2008). In our middle-aged population results, we observed that earning a high yearly job income was a top contributor to grey matter region volume in the dmPFC, vmPFC and AM, compared with the other examined traits. As a marker of socioeconomic affluence, the amount of yearly income a person earns aids in tracking one’s own place in the hierarchical layers of society as well as the social status of other individuals. A previous functional MRI study explored the neural processing of social hierarchies and found that participants generally tend to focus more on superior versus inferior individuals (Zink, Tong et al., 2008). In an unstable social hierarchy, brain regions including the AM and mPFC were recruited when a participant viewed a superior individual (Zink, Tong et al., 2008). The authors attribute their findings in the AM and mPFC to emotional processing and impression formation of other people in a social setting. Hence, our findings on income as a dominant trait manifestation in three social brain regions are consistent with the possibility that social status may have volume adaptations in the mPFC during midlife.

Furthermore, social status has consequences for and resonation with well-being across the lifespan (Sapolsky, 2005). Our examination of ~10,000 middle-aged individuals has singled out health satisfaction to yield a dominant trait effect in the vmPFC. Relatedly, a structural MRI study investigated the relationship between socioeconomic disparity and grey matter volume in ~450 middle-aged adults (Gianaros et al., 2017). The authors found that socioeconomic disadvantage, indexed by measures including household income and unemployment status, was linked to reduced grey matter in brain regions of the frontal and temporal lobes, including the orbitofrontal cortex. The authors suggest that socioeconomic disadvantage may show characteristic neural manifestations and ultimately lead to downstream implications on one’s health status (Gianaros et al., 2017). Together with our present population results, these considerations invite the speculation of mPFC and limbic involvement in maintaining social status, which may be protective of one’s health.

Being more of a morning person is associated with behavioral tendencies that promote prosociality, social connection and cooperation, such as conscientiousness and agreeableness (Tsaousis 2010; Takeuchi, Taki et al., 2014). In turn, conscientiousness has been linked to career success and overall physical and psychological well-being across the lifespan (Kern, Friedman et al. 2009; Lodi-Smith and Roberts 2012). Compared with the other examined traits, being a morning versus evening person showed a dominant trait association in the FP. Located in the mPFC, the FP is a region that is not only closely linked to perspective-taking (Lewis et al., 2011) but also future- and goal-oriented processes (Tsujimoto, Genovesio & Wise, 2011). In line with our social brain-trait associations, a previous voxel-based morphometry study found morning chronotype to be associated with grey matter volume in several brain regions, including the mPFC (Takeuchi, Taki et al., 2014). Moreover, a structural MRI study found higher levels of conscientiousness to relate to larger grey matter volume in mPFC regions (Jackson, Balota et al. 2011). Thus, having an early-riser biorhythm may perhaps provide far-reaching benefits, such as achieving certain career milestones. In the study from Takeuchi and colleagues, the authors found that morning types had higher levels in individual measures of prosociality and self-discipline compared to evening types, as well as lower levels of attentional or emotional problems (Takeuchi, Taki et al., 2014).

Relatedly, our population-level results revealed that the “early-bird” chronotype also emerged as a dominant trait association in the HC of the limbic social brain. The HC is also largely involved in memory processes, and has been thought to be involved in social monitoring, such as processes related to adhering to social norms (Goel & Dolan, 2007). Indeed, previous research has shown that being a morning versus an evening person is related to better adherence to social norms, self-control, cooperation, respecting authority and the social desire to give off a positive impression (Díaz-Morales, 2007). Additionally, being more of a morning person may sometimes require overriding one’s natural sleep-wake cycle to conform or adjust to the societal norm, even if one may be more of an evening person (Díaz-Morales, 2007). Conversely, the “night-owl” chronotype has been linked to measures related to risk-taking, creativity and resistance to acting in a traditional manner (Díaz-Morales, 2007). The conjunction of these previous behavioral findings and our present population-level results support the mPFC and its limbic connection to the HC to relate to morning versus evening orientation.

## Conclusion

By adopting a probabilistic generative framework at population scale, we jointly put to the test 40 traits with their correspondences in the social brain. This modelling strategy allowed isolating unique contributions of lifestyle traits that emerged as dominant in explaining imprints in brain structure. We confirm and extend previous findings by providing neurobiological evidence in 10,000 middle-aged individuals, which makes steps towards reintegrating three disparate fields of neuroscientific inquiry: social interaction, personality and demographics.

On the one hand, our quantitative findings can be interpreted to indicate that the decades-long accumulation of social skills and experience in societal roles may resonate in brain circuitry necessary for navigating social environments. On the other hand, as an alternative read-out, distinct individual, interpersonal and societal predispositions feed into life choices and thus life experience well into midlife. Put differently, our association study is impotent to disentangle the effects of nature versus nurture. However, our population-level evidence provides a glimpse into how different lifestyles may be reflected in long-term plasticity effects in brain circuitry.

